# Intra-V1 functional networks predict observed stimuli

**DOI:** 10.1101/2022.10.20.513108

**Authors:** Marlis Ontivero-Ortega, Jorge Iglesias-Fuster, Jhoanna Perez-Hidalgo, Daniele Marinazzo, Mitchell Valdes-Sosa, Pedro Valdes-Sosa

**Author notes:** Both contributed equally (Mitchell Valdes-Sosa, Pedro Valdes-Sosa).

## Abstract

Several studies suggest that the pattern of co-fluctuations of neural activity within V1 (measured with fMRI) changes with variations in attention/perceptual organization of observed stimuli. Here we used multivariate pattern analysis of intra-V1 correlation matrices to predict the level and shape of the observed Navon letters. We examined the inter-individual stability of network topologies and then tested if they contained intra-individual information about stimulus shape or level that was tolerant to changes in the irrelevant feature. The inter-individual classification was accurate for all specific level and letter-shape tests. These results indicate that the association of V1 topologies and perceptual states is stable across participants. Intra-participant cross-classification of level (ignoring shape) was accurate but failed for shape (ignoring level). Cross-classification of stimulus level was more accurate when the stimulus-evoked response was suppressed in the fMRI time series and not present for correlations based on raw time series, stimulus-evoked beta-series, or simulations of the effects of eye movements measured in a control group. Furthermore, cross-classification weight maps evinced asymmetries of link strengths across the visual field that mirrored perceptual asymmetries. We hypothesize that feedback about level information drives the intra-V1 networks based on fMRI background activity. These intra-V1 networks can shed light on the neural basis of attention and perceptual organization.

## Introduction

In brain network analysis, a single node frequently represents each cortical area. Hence, the functional magnetic resonance imaging (fMRI) time series from each region’s voxels (usually obtained during the resting state) are averaged (Schaefer et al., 2018; Sporns & Betzel, 2016). This summarization overlooks signs of neural cooperation-indexed by co-fluctuations of activity-within cortical regions. Furthermore, resting state (as opposed to task-fMRI) connectivity is not optimal for probing specific cognitive functions (Finn, 2021). Thus, more intra-region network studies using task-fMRI are needed.

The internal connectivity of the primary visual cortex (V1) is a promising candidate for inquiry since much is known about this area’s internal specialization, response to task requirements, and neuronal subtypes, which could aid the interpretation of task-fMRI studies (Saenz & Fine, 2010; Seidemann & Geisler, 2018). The close correspondence between V1 macroscopic anatomy and retinotopic organization (Benson et al., 2012; Benson & Winawer, 2018) simplifies across-individual comparisons: there is a consistent mapping of the visual field (VF) mapping onto V1. Moreover, V1 has small population receptive fields in V1 (pRFs, Wandell & Winawer, 2015). Therefore, the synchronization of fMRI activity between cortical sites that are sufficiently separated (therefore mapping points distant in the VF) suggests one (or both) of two mechanisms: 1) lateral interactions within V1 (Liang et al., 2017); 2) feedback from high-order visual areas (Lamme & Roelfsema, 2000; Liang et al., 2017).

Two studies have shown that fMRI activity from different V1 sites is more strongly correlated when mapping parts of the same, as opposed to distinct, visual objects (Nasr et al., 2021; Valdes-Sosa et al., 2022). These studies used within-individual univariate statistical testing on measures from a few V1 ROIs (Nasr et al., 2021) or many between-site edges (Valdes-Sosa et al., 2022). However, univariate testing ignores systematic associations between features, possibly missing more complex patterns (Davis & Poldrack, 2013; Haynes & Rees, 2006). Also, it does not address the stability of network topologies across individuals, which is important when small numbers of participants are studied (Calhoun, 2022).

Here, we asked if the pattern of intra-V1 connectivity - measured by fMRI activity-allows prediction of the distribution of attention. More specifically, we tested if it was possible to predict with multivariate pattern analyses (MVPA) which level of a compound figure (global or local) the participants were paying attention to. MVPAs were applied to the fMRI connectivity matrices from the Valdes-Sosa et al. study. That study used modified Navon figures (Iglesias-Fuster et al., 2014) that temporally segregated the display of global and local letters within the same matrix of lines. Four stimuli, global and local ‘E’s and ‘U’s, were presented. Inter-individual MVPA (Wang et al., 2020) was used to assess the stability of network topology associated with each type of stimulus across individuals (classifiers trained with data from n-1 individuals of individuals were tested on data from the excluded individual). Inter-individual MVPA suffers when the spatial registration of fMRI patterns is inaccurate, a problem that is fortunately mitigated for V1 data given the excellent between-subject registration of retinotopic mappings mentioned before. With this approach, we tested “specific” classifications for each feature: Level and Letter identity (or shape). We called them “specific” because the irrelevant feature did not change.

Conversely, within-individual cross-classification MVPA (Kaplan et al., 2015) was used to probe the nature of the associations between more abstract stimulus attributes (level and letter identity) with the V1-connectivity patterns. Cross-classifications can show if intra-V1 network topologies are distinctive for each level irrespective of letter shape or letter shape irrespective of level. The success of this learning transfer in cross-classification implies that a primary attribute’s neural representations are “tolerant” to changes in a secondary attribute. Tolerant and specific classifications probably are due to distinct neural circuitry.

When studying the correlation of fMRI activity within V1, controlling for between-site spurious correlations induced by the stimulus-evoked responses is necessary (especially if these confounders differ between conditions). The stimuli may activate distinct - noninteracting-cortical sites in parallel. In addition to experimenter-controlled stimulus change, gaze displacements over the stimuli could modify effective retinal stimulation, abruptly presenting different stimulus edges to V1 sites with each fixation. A previous fMRI study of Navon stimuli found diverging eye-movement patterns when viewing global and local stimuli (Sasaki, 2001). Therefore, the bursts of fMRI activity triggered by the fixations on different parts stimuli could also generate intra-V1correlations that may differ between conditions.

We tackled here the issue of spurious stimulus-or fixation-induced correlations with two approaches. First, we compared MVPA classification accuracy for different stimulus-evoked response/background-activity ratios. This ratio can be modified during the fMRI preprocessing steps before calculating the correlations (Al-aidroos et al., 2012). If MVPA classification accuracy improves (or is unaltered) after evoked-response component reduction, discrimination is unlikely due to spurious stimulus-induced correlations. Second, we used a recent image-computable model based on Gabor pyramids (Kay et al., 2013) that predicts BOLD responses to known stimuli in the early visual cortices. We simulated the possible effects of eye movements (with gaze tracking records measured in an offline experiment) on each simulated time series.

Additionally, the retinotopic organization of V1 allows meaningful interpretation in the visual field of the “forward models” (Haufe et al., 2014) of the feature weight maps underlying successful classifiers. V1 network edges connect cortical sites for which population receptive fields (pRFs) have been mapped (Wandell & Winawer, 2011, 2015). The weight of each link can thus be plotted in the visual field as a line connecting the positions (defined by the pRFs) corresponding to two nodes. The network edges contributing most to the predictions should have a characteristic spatial pattern corresponding to known psychophysical asymmetries within the visual field (Levine & McAnany, 2005; Nasr et al., 2021).

## Materials and Methods

The fMRI data used here have been previously described in other publications. A summary description is provided below; see Valdés-Sosa et al., 2020 and Valdés-Sosa et al., 2022, for a full explanation.

### Participants

Twenty-six human volunteers (ages 23 to 28 years; 9 females) participated in the study. All had normal, or corrected-to-normal, vision, did not present any medical condition and were right-handed except for two cases. The procedures were approved by the ethics committees of the University for Electronic Science and Technology of China (UESTC) and the Cuban Center for Neuroscience. Participants gave written informed consent in compliance with the Helsinki Declaration.

### Stimuli and tasks

Modified Navon figures (Figure 1) were presented in the experiment. Two letters, E and U, were presented at both levels in a blocked stimulus paradigm. The 1 s letter presentation alternated with a background mask also flashed for 1 s. The letters were made from white lines on a black background with about 2.0 s of visual angle (DVA) width and 5.3 DVA height. This matrix was built out of smaller placeholder elements shaped like ‘8’s (with visual angles about 40 minutes of DVA wide and 1 DVA and 3 minutes high). Only one letter was shown in each block, in which participants were required to report the number of minor deviations in letter shape. The stimuli were projected on a screen at the subject’s feet, viewed through an angled mirror fixed to the MRI head-coil, and were generated using the Cogent Matlab toolbox (http://www.vislab.ucl.ac.uk/cogent.php).

**Figure 1.**
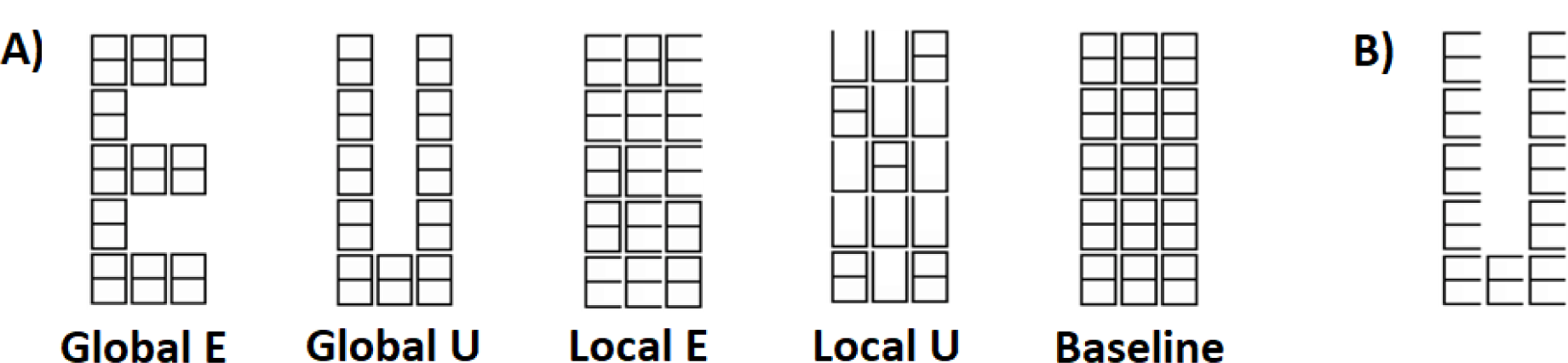
Navon figures. A) Modified figures used in the experiment. Two letters (E and U), and two level (Global and Local). B) Example of traditional Navon figure.

Blocks had 44 sec of duration and consisted of an initial cue (’Global’ or ‘Local’) (1 s), followed by a 19-sec baseline, 20 s of the same letter (1-sec repetition) and were ended with a 4 sec wait period where the number of shape deviations was reported. Five runs were presented in 24 participants and four runs in two, each consisting of 12 blocks (3 blocks for each letter: EG, EL, UG, and UL).

### Data acquisition and Image pre-processing for the main experiment

Recordings were carried out with a GE Discovery MR750 3T scanner (General Electric Medical Systems, Milwaukee, WI, USA) using an eight-channel receiver head coil. Functional images were obtained with a T2*-weighted echo planar imaging sequence (TR=2.5s; TE=40 ms; flip angle=90^○^) with a spatial resolution of 1.875 x 1.875 x 2.9 and 135 images per run. A T1-weighted image was also obtained with 1 x 1 x 0.5 mm resolution.

Initial pre-preprocessing of functional data included discarding the first five volumes of fMRI in all runs, artifact correction (using ArtRepair toolbox (http://cibsr.stanford.edu/tools/ArtRepair/ArtRepair.htm), followed by slice-timing, head motion correction (with the extraction of motion parameters) and unwarping with SPM8 (http://www.fil.ion.ucl.ac.uk/spm/). The T1 Image was segmented and normalized to MNI space using SPM12 to extract nuisance parameters from fMRI activity in white matter (WM), and cerebrospinal fluid (CSF) that were included in the general linear model described below. For each subject, these masks were created using a threshold of tissue probability greater than 0.9. The CSF mask was also restricted to the ventricles using a template in MNI space (https://sites.google.com/site/mrilateralventricle/template).

Cortical surfaces (white and pial) were reconstructed from the T1 Image for each subject using Freesurfer (http://surfer.nmr.mgh.harvard.edu), registered to the FsAverage template, and subsampled to 81924 vertices. The mid-gray cortical surface was co-registered with the functional data. Then the fMRI time series were interpolated to each mid-gray cortical surface and high-pass filtered with a time constant of 128 s. Also, for subsequent spatial smoothing of the functional data in V1, discs with 5mm radii were defined over the FsAverage surface using the Surfing toolbox (http://surfing.sourceforget.net).

### Estimation of fMRI background activity connectivity matrices

Background activity (BA) was defined as the residual time series of each surface vertex after regressing out the effects of the stimuli (evoked response) and 64 nuisance parameters (a standard set of nuisance variables to eliminate the effects of noise, artifacts, and physiological contaminants) applying a general linear model (GLM). The nuisance regressors included the primary motion parameters, their derivatives, and the quadratics of both these sets (24 motion regressors in total). Physiologic noise was modeled using the aCompCor method (Behzadi et al., 2007) on the time series extracted separately from the masks of WM and CSF in ventricles in volume space. The first five principal components from each set of time series, the derivatives of these components, and the quadratics of all these parameters were obtained (40 regressors in total).

### Preprocessing schemes

Two variants of the GLM were used. In the main analysis, each stimulation block was modeled as a square wave convolved with the canonical hemodynamic function from SPM 12 (c-HRF). In another case (RAW), the stimulus effect was omitted from the design matrix..

The residual time series in V1 vertices, obtained after GLM, were smoothed by averaging with the time series of its neighbors in the 5 mm discs mentioned above. The studied V1 region was restricted to the central 8 DVA of eccentricity in a probabilistic a priori map (Benson & Winawer, 2018). Next, the time series were segmented into blocks (considering the time shift introduced by the hemodynamic function). These segments were linearly detrended, and segments corresponding to the same stimulus were concatenated (120 points-equivalent to 300 seconds-for each stimulus in 22 participants and about 96 points – equivalent to 237.5 seconds-in 4 subjects). We base this procedure on prior reports that resting state fMRI analyses with concatenated data were not significantly different from the analyses of continuous data in multiple aspects (Cho et al., 2021; Zhu et al., 2017).

Finally, intra-V1 connectivity matrices were estimated by calculating the Pearson correlation coefficient between the pre-processed time series for each vertex of V1 and segregated by stimulus condition (EG, EL, UG, UL) in all participants. For the classification analysis, all matrices were vectorized. The correlation values were converted to z-values using the Fisher transformation, negative values were set to zero, and missing values in each subject’s vector were substituted by the median value of the other participants without missing values. These missing values could have been due to noise or BOLD signal dropout at specific cortical vertices. However, they were few and concentrated in restricted areas of the visual field (see Valdes-Sosa et al., 2022).

### Two types of Classification analysis

Intersubject MVPA (to assess stability across participants) and within-individual cross-classification MVPA (to assess discrimination invariance) were performed, in which the possibility of predicting observed stimuli from the intra-V1 connectivity matrices was measured. In all tests, the connection strengths between all node pairs were used as features, and a support vector machine (SVM) was used as the classifier (with the default parameter C=1). Feature selection was performed to eliminate some irrelevant connections by applying a two-tailed t-test between conditions on the training data and retaining links with significant t-values (p <0.01). Prediction accuracy (Acc) and the area under the ROC curve (AUC) were used to assess the performance of each classifier. The statistical significance of deviation from a random classification (0.5 for Acc and AUC) was estimated by permutation testing (1000 times), in which stimulus labels were randomly changed (Valente et al., 2021).

### Intersubject-classification tests

Specific inter-individual classification tests were performed to evaluate if the pattern of association between stimulus conditions and V1 network topology was stable across participants. These tests were carried out with cross-validation in a leave-one-subject-out (LOSO). Thus, training was based on the data of n-1 participants and testing on the data of the remaining participant. The four discriminations tested were level (Global vs Local), separately for the ‘E’ and the ‘U’ stimuli, and letter shape (’E’ vs ‘U’), separately for the Global and the Local stimuli. Each iteration of the LOSO consisted of 50 training samples and two testing samples.

### Intraindividual cross-classification tests

Cross-classification tests were employed to see if models built for a relevant feature (e.g., level) were invariant to changes in another irrelevant feature (e.g., letter identity). The data from all subjects were divided into two sets of pairs (each with 52 observations) to test the invariance of level discrimination with respect to letter identity: those associated with EG-EL, and with UG-UL. The data was also divided into matrices associated with EG-UG and EL-UL to test the invariance of letter discrimination with respect to level. The classifier was trained alternating which pair was used for training and which for testing, and the two accuracies and AUCs were averaged and reported. Thus, two instances of the cross-classifier were always calculated (e.g., level for ‘E’ and ‘U’). Note that cross-validation was unnecessary in these tests since the classifier was trained with data from one pair of conditions and tested with independent data related to the other pair. Permutation tests were based on randomizing the labels of the training data in both directions of the test. These tests with repeated with beta-series correlations; in other words, with the variations in beta weights for each stimulus over time at each V1 cortical node.

### Comparison of classification after different fMRI preprocessing schemes

To determine the relative contribution of stimulus-evoked activity, compared with background activity, we compared classification accuracy after different fMRI pre-processing schemes. One cannot assume that the GLM completely eliminates the evoked activity since HRF modeling is likely imperfect. But if classification is not affected, or improves, after the reduction of the evoked response via GLM, background activity must play an important role in the classifications. Thus, we measured the accuracy of classifiers when using connectivity matrices obtained with the RAW data (stimulus effects untouched) in addition to the main analysis with c-HRF deconvolution (deconvol HRF). Both specific classifications, as well as cross-classifications, were examined. The data were bootstrapped over participants (100 times), and the results were employed to build the confidence intervals for classification accuracy.

### Control eye movement experiment

A control experiment was conducted with 19 additional participants (13 female y 6 male, age range 21-61, median = 42), all Cuban university students or graduates. All had normal (or corrected to normal vision), no history of neuropsychiatric diseases, and 16 were right-handed. Eye movements were measured while the subjects observed the same stimuli - and performed the same task-from the fMRI experiment but with slightly larger stimuli. A 34 x 27 cm monitor screen (1280 × 1024 pixels resolution) was used. A chin and forehead rest fixed the participant’s head position at 69 cm from the screen; therefore, the stimuli were about 7.16° wide and 2.9° high.

We used an EyeLink® 1000 Plus Version 1.0.6 Desktop Mount system (SR Research Ltd., Ontario, Canada), to measure eye position by recording corneal reflection and dark pupil with a video-based infrared camera and reflective mirror. These measurements had a spatial resolution of 0.01° of visual angle and a temporal resolution of 1000 Hz. The viewing was binocular, but the recording was monocular. Calibration and validation of the measurements were performed before each experimental session. The fixations during the 20 s stimulation blocks were separated into sets corresponding to the four stimulus types.

The distribution of eye movements needs to be distinct across different stimuli, and to be consistent across participants, for the former to influence the between-subject identification of presented stimuli from their associated intra-V1 networks. Linear mixed models (LMM) were applied to analyze the deviation of gaze position from the central fixation point, and fixation duration, as a function of Level (Global Vs. Local), Letter (’E’ vs. ‘U’) and their interaction, in addition to trial order within stimulus category (from 1 to 5), based on the following formula (in Wilkinson notation):

*Disfix* ∼ 1 + Level + Letter + TrialOrder + Level: Letter + (1|participant)

*FixDur* ∼ 1 + Level + Letter + TrialOrder + Level: Letter + (1|participant)

V1 networks predict observed stimuli
where TrialOrder, is the order of presentation for each type of stimulus, *DistFix* is the distance of the gaze in pixels from the fixation, *FixDur* the duration of the fixations.

Eye movement data were analyzed using the program iMap14 (Lao et al., 2017). Fixation durations for each trial were projected back into their x and y pixel coordinates in the visual field as Dirac deltas of the appropriate amplitude. This fixation duration map was convolved with a two-dimensional Gaussian function with a10 pixel standard deviation, producing gaze heatmaps that were then downsized in pixels with a scale of 0.25. The resulting 3D matrix (trials x 320 x 256) Y was modeled in a mass univariate LMM as the dependent variable according to the following equation:

*FixY (x,y)* ∼ 1 +Level + Letter + TrialOrder + Level: Letter + (1|participant)

This and other LMM were fit by maximal likelihood (ML) using the fitlme function from the Statistics Toolbox™, Matlab 20122b. Original parametric statistical values were thresholded at a given p-value (0.05/the number of pixels). The cluster mass was obtained by summing the statistic values within each cluster and were later compared with a bootstrap distribution under the null hypothesis (1000 times) values allowing to correct the p values for the multiple comparisons inherent to the mass univariate analysis of heatmaps (here 81920 comparisons).

### Simulation of effect of eye movement in fMRI correlation matrices

To assess the potential influence of eye movements on the patterns of intra-V1 fMRI connectivity from our experiment, simulated time series were generated reflecting the estimated effects of gaze change on V1 voxel activity. As in previous work (Kay et al., 2008), V1 was modelled as a pyramid of Gabor filters. The bank of filters consisted of five resolution levels (1,2,4,8,16 cycles/FOV), eight orientations (0, 22.5, 45, 67.5, 90,112.5, 135, and 157.5°), and two phases which tiled the screen at evenly spaced positions according to the resolution (2,8,16,64,256 respectively). The stimulus patterns on the screen were convolved with the gaze trajectories for their corresponding blocks for each participant. The two phases and eight orientations from the output from the Gabor filter bank were collapsed at each position for all scales for each fixation episode. The simulated fMRI time series consisted of segments in which the filter output for each fixation was expanded in time according to their durations. The time series were convolved with the canonical HRF function and downsampled to the fMRI TR (2.5 s). The correlation matrices (size: number of Gabor-filters x number of Gabor-filters) of these time series were calculated for each stimulus in all participants. The same classification procedures used for the fMRI data were applied to these virtual correlation matrices. The outcomes were bootstrapped 100 times to obtain robust estimates of the central tendency (median) and dispersion (central 68% percentile range) for accuracy and AUC.

### Analysis of weight maps in the cross-classification tests

The weight maps of the SVM cross-classifiers were examined to determine which regions of V1 contributed most to classification tests. This analysis was not carried out for the specific intersubject tests since their weight maps are necessarily different, reflecting the retinotopic stimulation pattern. In contrast, the invariance of weight maps for one attribute despite changes in another implies a generalization beyond the precise pattern of retinotopic stimulation.

The stability of the weight maps was estimated by bootstrapping (1000 times) procedure. In each bootstrap iteration, new data was created by resampling with replacement across participants. The SVM was re-trained in each replication to obtain multiple models (for both instances of the irrelevant attribute). The estimated model coefficients were transformed into forward models as recently proposed (Haufe et al., 2014) producing “activation” maps (or HTWs) to enhance interpretability.

We then calculated the concordance of HTW map feature rankings between the two instances of the invariant cross-classifications. If the classifiers trained separately for two instances yield similar weight maps, then the features guiding the generalization of learning across the two situations are equivalent. Concordance was measured between the two instances for each bootstrap replicate using Kendall’s W index (which ranges from 0 for no concordance to 1 for perfect concordance). If the classifiers contain invariant information (i.e. tolerance to irrelevant feature change), the ranking of activations in the two HTW maps should be highly concordant. The 95 % bias-corrected and accelerated percentile confidence intervals of W were calculated by bootstrapping.

The HTW maps from the two cross-classification instances were averaged to enhance features that were relevant in both. This average HTW map was transformed into a robust pseudo-zscore (https://github.com/cvnlab/knkutils), obtained by dividing the median of the feature by one-half the width of the central 68% range (an analog of a standard error) of the bootstrap values. This Z-transformed HTW map was split into positive and negative portions (e.g., each favoring one level). The absolute values of these maps were fit with two models: a Gaussian distribution and the mixture of a null Gaussian and gamma distribution. The model with the lowest Bayesian information criterion (BIC) was used to estimate the respective false discovery rates (q=0.00001). Finally, the pseudo-zscores (representing edges in a connectivity matrix) surviving the FDR threshold were projected onto the visual field and shown as graph plots in visual field coordinates, using the mean polar angles and eccentricities for V1 vertices extracted from a population retinotopy prior map (Benson & Winawer., 2018).

We also calculated the 95 % percentile confidence intervals for each feature in the average HTW map. Significant features (links for which the confidence interval did not include zero) were plotted within the original correlation matrix space. Median HTW values were projected as graph plots onto the visual field as indicated above.

### Results Classification results in the main experiment

The results of accuracy and AUC for all classification tests are shown in Figure 2. The inter-subject MVPAs accurately decoded all specific instances of the level discriminations (Acc: 0.79, AUC: 0.89 for ‘E’ and Acc: 0.75, AUC: 0.85 for ‘U’) and of the letter discriminations (Acc: 0.73, AUC: 0.83 for Global and Acc: 0.69, AUC: 0.74 for Local), with all permutation tests highly significant. Thus, significant inter-subject classification showed that the V1 topologies associated with each stimulus condition were stable across subjects.

**Figure 2.**
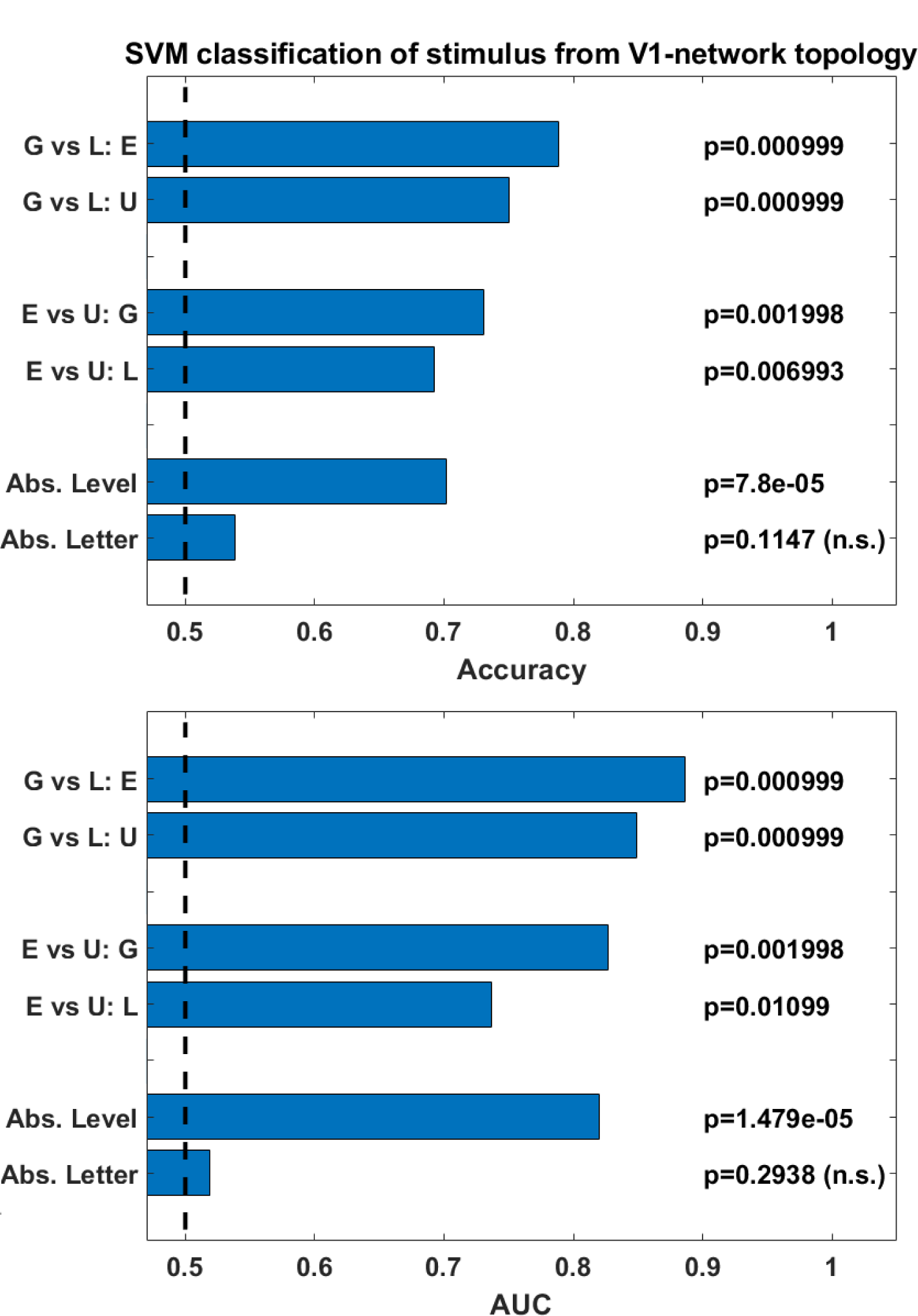
Classifier performance. The upper panel shows accuracy values in the bars. The lower panel shows AUC values in the bars. A vertical dotted line represents the chance level (0.5). Probabilities values from the permutation tests are shown in the numerical insets. G vs. L: E is the specific level classification with the letter “E,” and G vs. L: U is the specific case with “U.” E vs U: G is the specific letter classification at the global level, whereas E vs. U: G is the case for the local level. For the abstract cross-classifications (Abs.), the accuracies in two directions were averaged, and the p values were combined with Fisher’s method.

Also, invariant cross-classification for Level was highly significant (Acc: 0.70, AUC: 0.82), which means that a change letter identity did not adversely affect the discrimination. In contrast, invariant letter cross-classification was at almost chance level (Acc: 0.54, AUC: 0.52) and thus not significant in the permutation test. These results indicate that the spatial scale of stimuli (invariant to shape) is well reflected in the topology of intra-V1 networks, whereas letter shape independent from the spatial scale is not.

An analysis with correlation matrices based on beta-series (reflecting the changes over time of fMRI activations did not yield significant results for the cross-classifications.

### Classification results using different fMRI preprocessing schemes

A comparison of classification accuracies resulting from the two preprocessing strategies is shown in Figure 3. For both schemes (RAW and deconvol HRF), specific classification accuracies were well above chance. This was also true for the connectivity matrices calculated from the beta-series. All cross-classifications for RAW and beta-series connectivity matrices were at the chance level. This was also true for cross-classifications of letter identity across levels, which were at the chance level. Abstract level discrimination (i.e., cross-classifying level across letter identities), was significant only for both c-HRF preprocessing.

**Figure 3.**
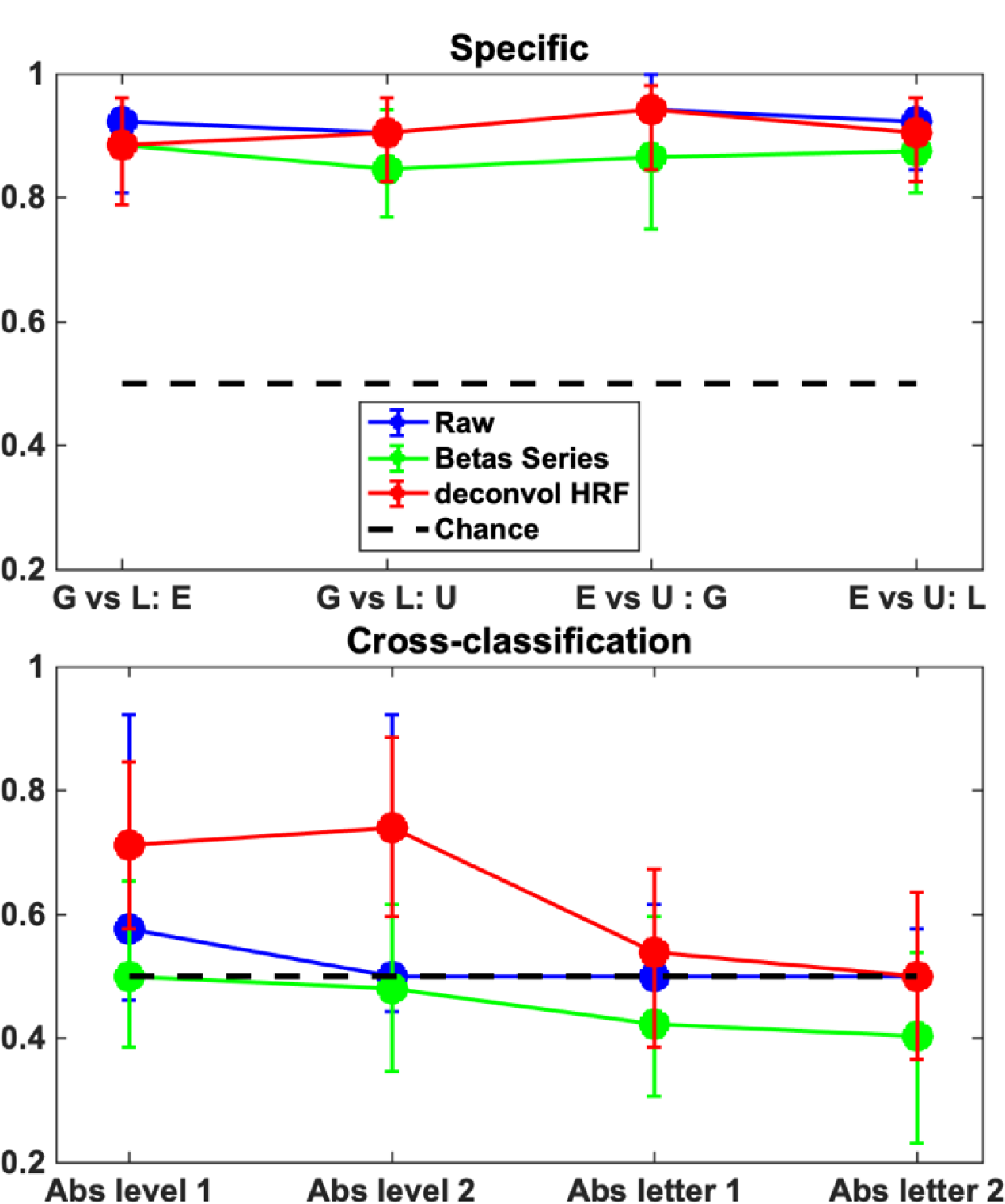
Comparison of bootstrapped classification accuracies under the two pre-processing strategies and for the beta-series. The markers represent the median and the whiskers, the 95% confidence interval of the bootstrapped values. Abbreviations are the same as in Figure 2. However, in this figure, the two cross-classification directions are not averaged.

### Comparison of eye movement across stimuli

There was no effect of Letter, and only a marginal effect of Level (t(1580) = 1.92, p < 0.06 uncorrected) on the mean distance of fixations from the center of the screen, with a beta value of only about 20 pixels (0.4°) in size. The interaction of Letter and level was significant (t(1580) = 5.65, p < 0.001) with a beta of about 14 pixels (0.3°). The effects of Level (t(1580) = 3.09, p < 0.001) and Letter (t(1580) = 3.28, p < 0.001) on fixation duration were highly significant, but this was not so for their interaction. Fixations were about 169 ms longer for Local than Global stimuli and 223 ms for the ‘U’ than the ‘E’. The effect of trial order was also significant (t(1580) = 4.73, p < 0.001), with a decrease of about 41.4 ms for each repetition of a stimulus type.

Gaze heat maps are critical here since they reflect cumulative and interactive effects of fixation position and duration that could potentially affect the fMRI time series. Neither Letter nor Level was significant after correction for multiple comparisons. Even with an uncorrected p-value of 0.05, was any effect found. Heatmaps for the four stimuli used in the experiment were very similar (Figure 4).

**Figure 4.**
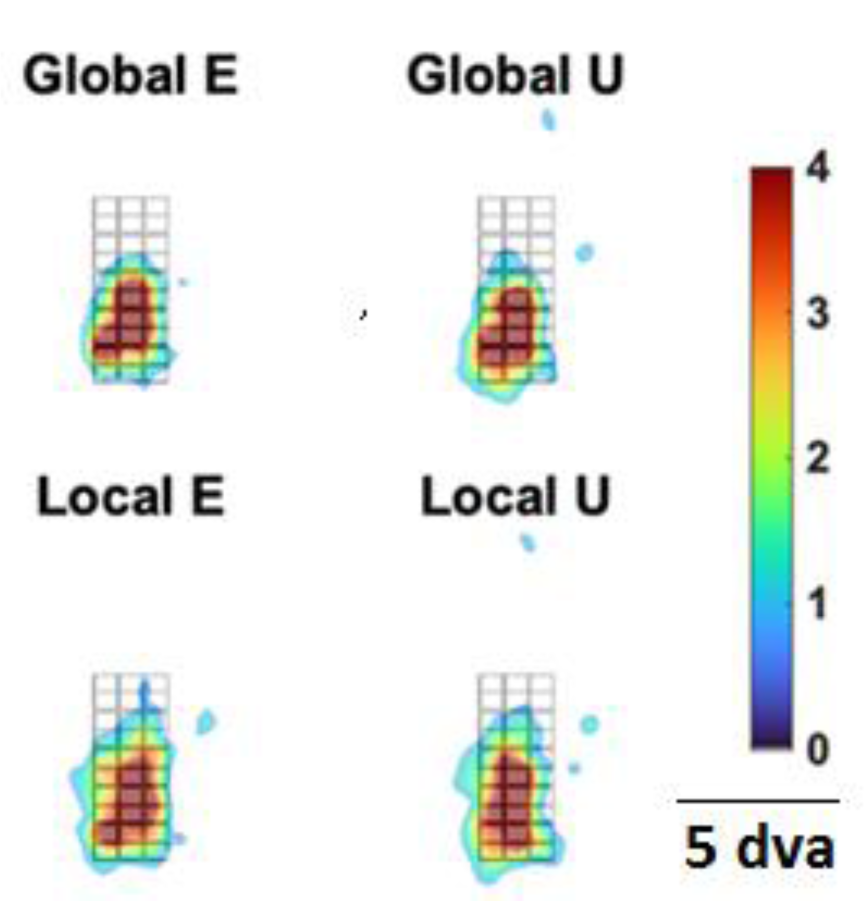
Mean heatmaps of gaze distribution for the four conditions. The contour images (using an arbitrary common scale for the four stimuli) are superimposed on the base mask stimulus. The vertical line corresponds to 5 DVA.

### Classification of simulated correlation matrices

The median (and 68% interquartile range) of classification accuracy and AUC values for the both cross-classifications (Level and Letter) were at chance level, as shown in Figure 5. Thus, neither the abstract level (independent from the letter shape) nor the abstract letter shape was decoded accurately from the simulated correlation matrices.

**Figure 5.**
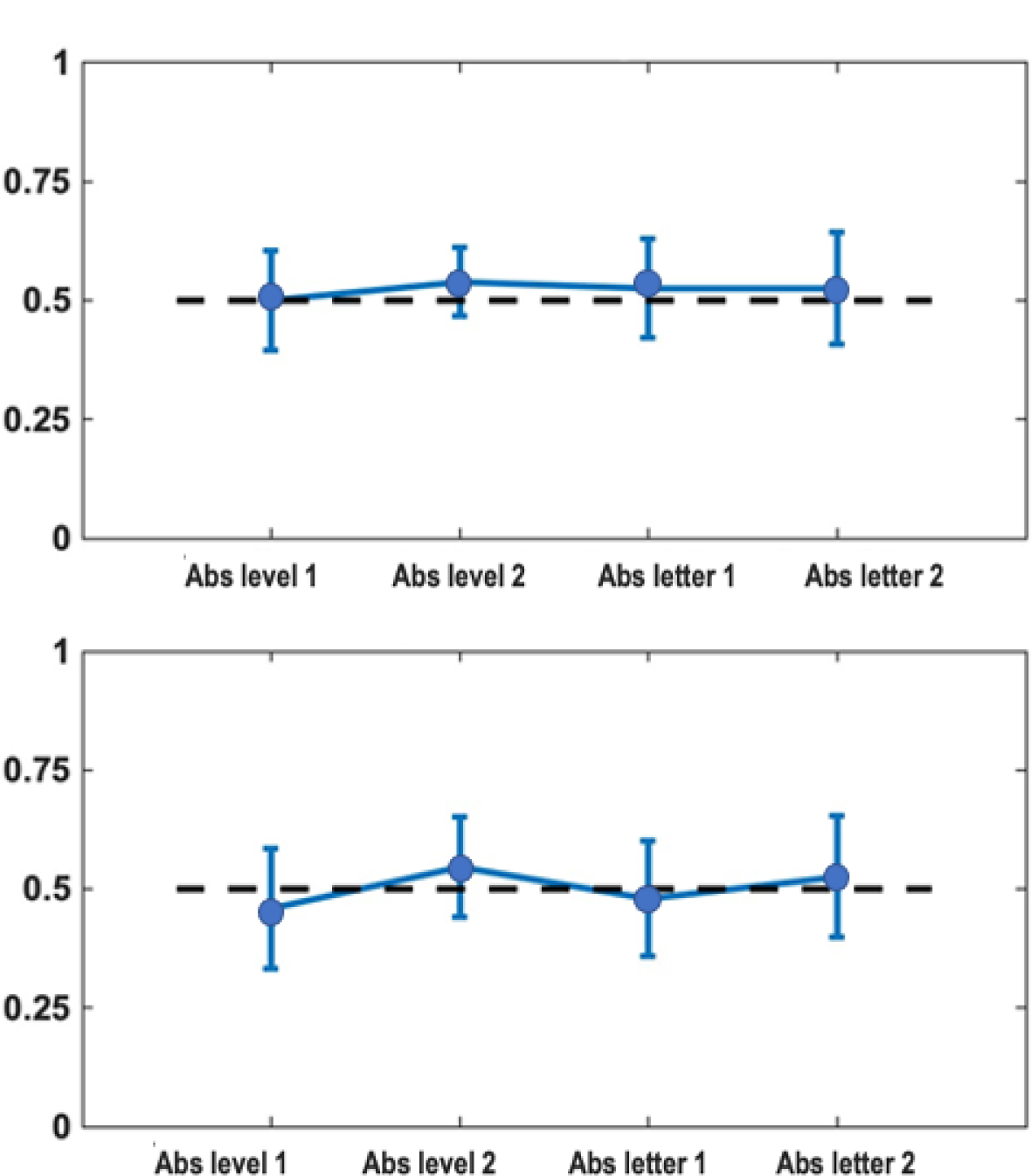
Performance for the cross-classification tests (bootstrap across participants). In the top panel, the proportion of accurate responses. In the bottom panel, the AUC values. The discontinuous black line represents the chance level. In all cases, the classifier was trained on discriminating one feature and then tested with simulated data after changes in the non-trained feature.

Contrariwise, the classification of simple, specific, discriminations was (Figure 6) significantly above the chance level for all examples.

**Figure 6.**
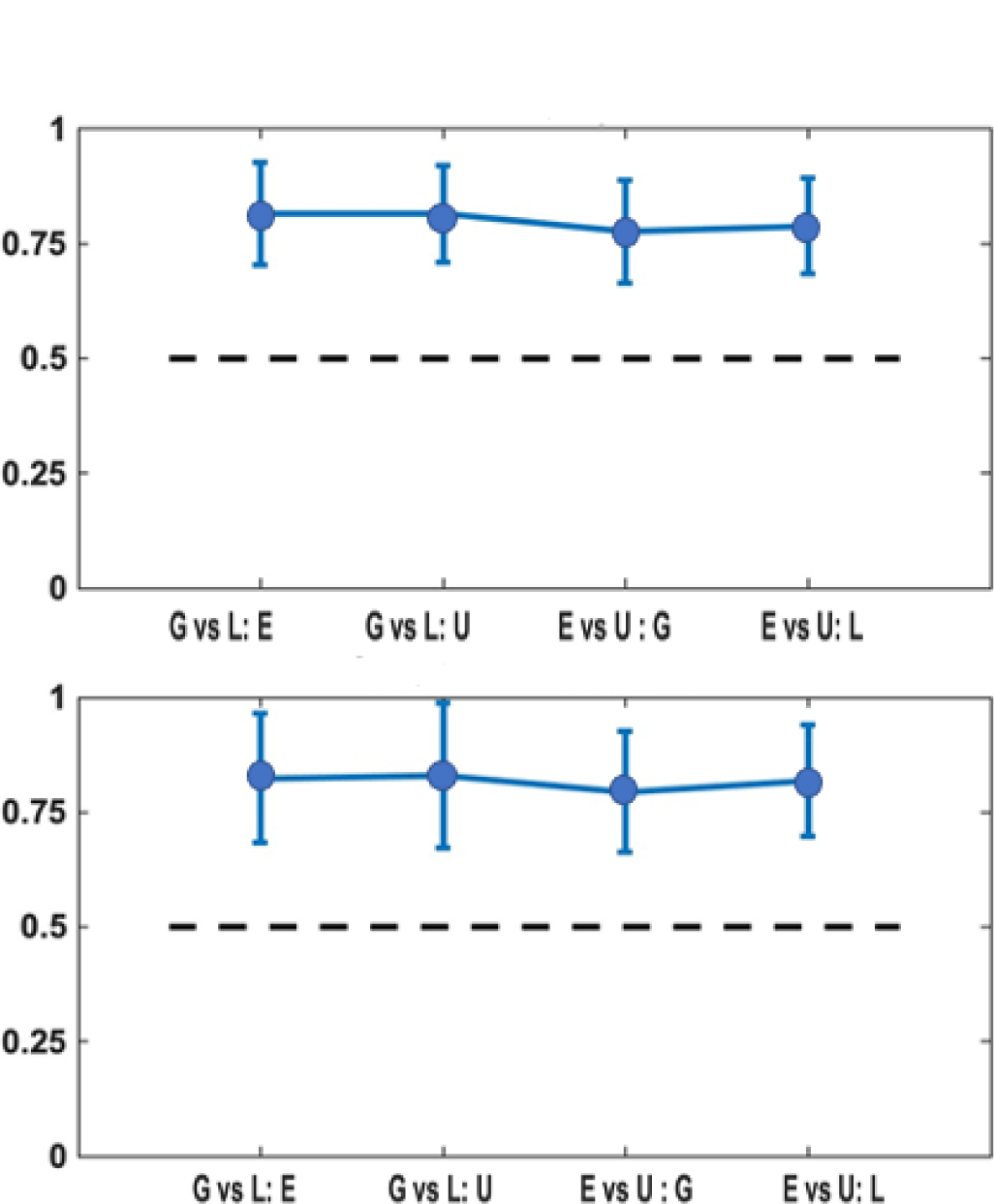
Performance for the simple classification tests (bootstrap across participants). Same conventions as in Figure 2.

### Weight maps for cross-classifications

Since accuracy in the cross-classifier for letter identity was at chance, we only show HTW results for level. A comparison of the HTW maps for the classifiers discriminating the Global/Local ‘E’ and for the Global/Local letter ‘U’ shows they were very similar. The observed W values between the two within-individual SVM classifiers for level was 0.77, with a 95% BCa confidence interval [0.65-0.83], which was well above zero (see the histogram in Figure 7). This result is congruent with accurate learning transfers between the two instances of level cross-classifiers.

**Figure 7.**
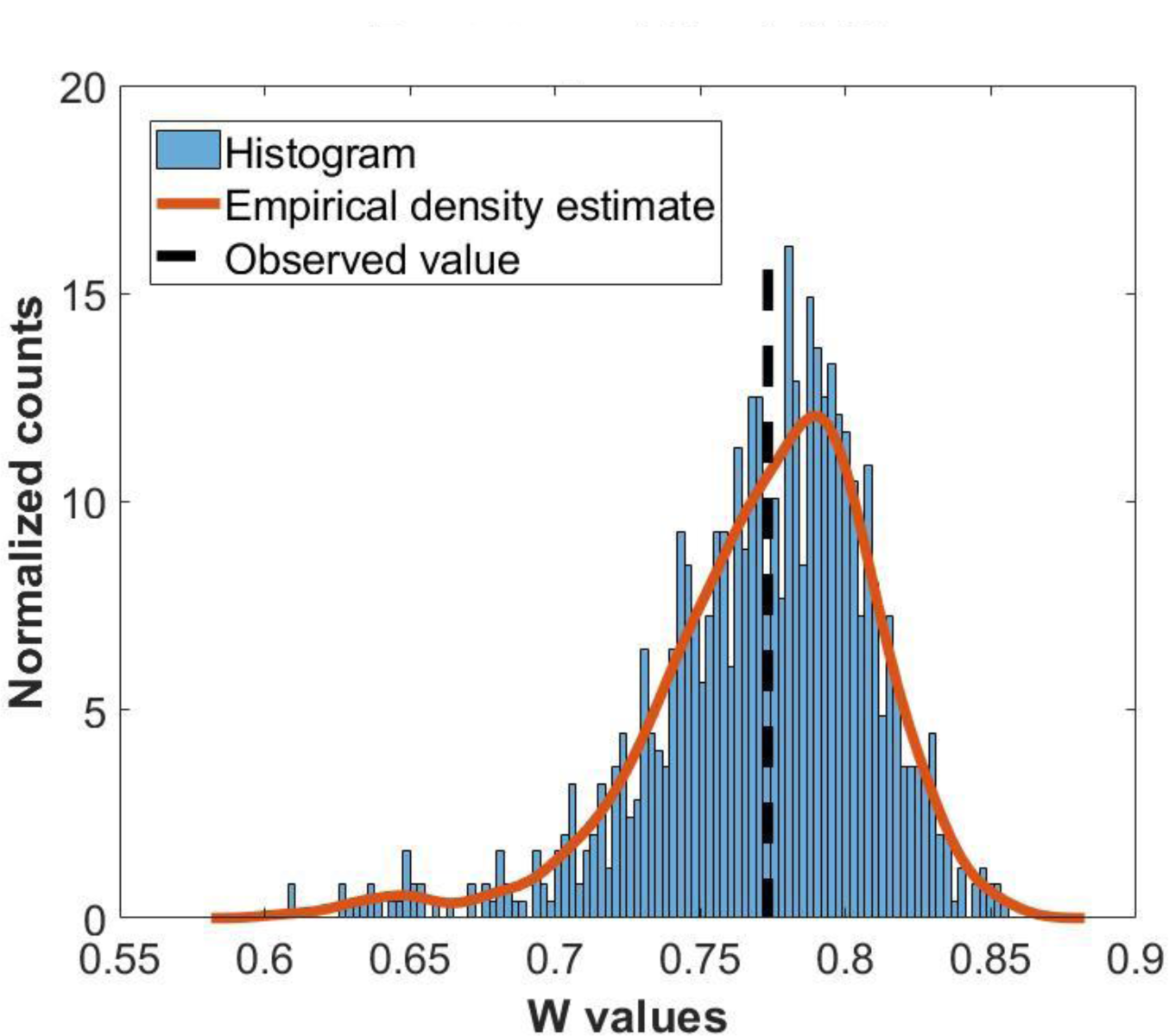
Distribution of bootstrapped Kendall W values. The resampling was over the 26 participants.

Maps of the link pseudo-z scores on the SVM Haufe-transformed weights (HTW) for the invariant level classifiers are plotted in the visual field in Figure 8. The Gaussian + Gamma mix provided better fits (lower BIC) than the lone gaussian for both positive and negative HTWs. The map for negative HTWs (favoring Global), after thresholding with the FDR calculated from the distribution mixture fit, exhibits many links crossing the vertical meridian, mainly in the lower visual field. The map for positive HTWs (favoring Local) shows many links circumscribed to the lower right and left visual quadrants.

**Figure 8.**
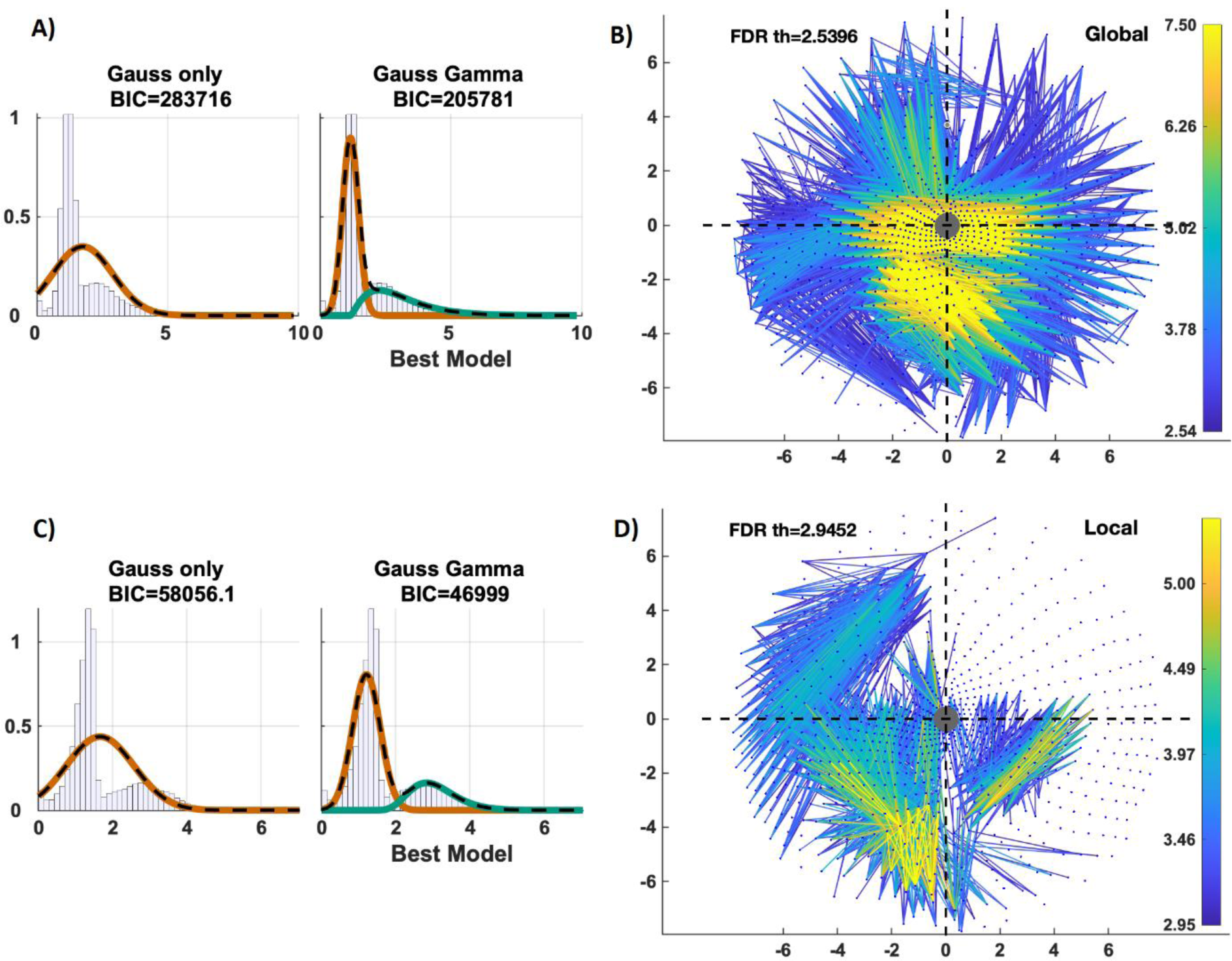
Analysis of standardized SVM Haufe-transformed weight (HTW) maps for the inavariant level classifiers. The top row (A, B) corresponds to negative HTWs (indicating Global, although absolute values are shown), and the bottom row (C, D) corresponds to positive HTWs (indicating Local). The estimated distribution mixtures are shown on the left side of the figure, with data histograms in black, the Gaussian component in red, and the gamma component in green. On the right side, graph plots in the visual field of the features surviving the FDR threshold, with color representing the magnitude of the pseudo-zscores. The gray circle represents the center of gaze, x and y axis represent DVA. Each light-blue dot corresponds to the center of a V1 pRF in the visual field.

Congruent results were obtained for the confidence interval analysis based on the HTW bootstrap. The links with confidence intervals that did not include zero are shown in Figure 9 (corresponding to values above chance). In this correlation matrix space, interhemispheric links dominate in the Global condition (occupying the block anti-diagonal), whereas intrahemispheric links dominate in the Local condition (occupying the block anti-diagonal). When projected onto the visual space, the median values of significant HWT values tend to cross the vertical meridian of the VF in the lower quadrants for the Global condition. Significant HWT values in the Local condition tend to stay within single visual field quadrants, especially the lower ones. These results are largely congruent with the findings in Figure 8.

**Figure 9.**
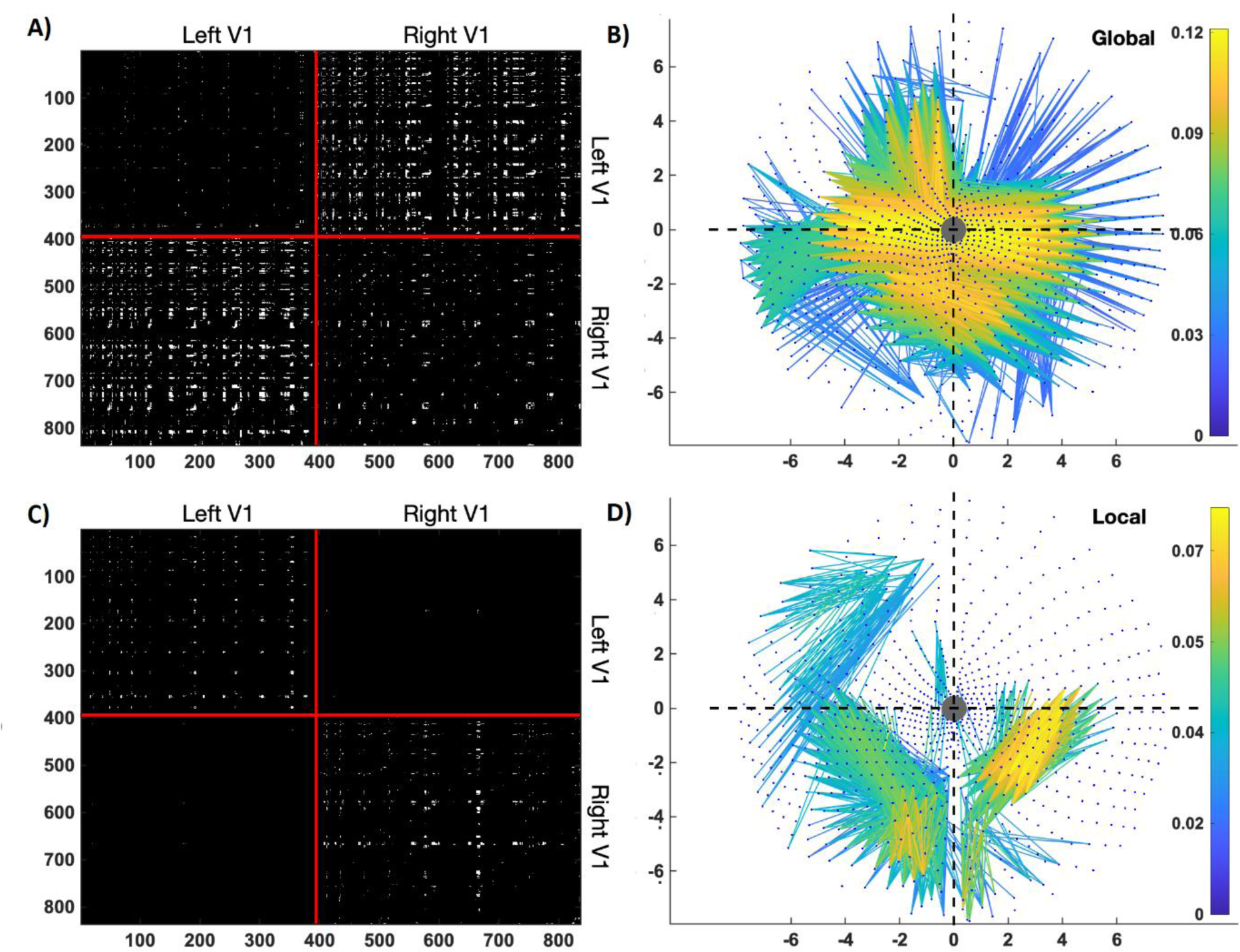
Confidence Intervals (95%) analyses of Haufe-transformed weight maps for the invariant level classifier. Cases in which the intervals did not contain zero were selected and plotted separately for positive activations (top row: A, B) and negative activations (bottom row: C, D). On the left, the surviving features are marked in white within the original V1 vertex correlation matrices. The numbers on the axis indicate the position in an arbitrary list of V1 cortical vertices. The red lines indicate where the list of vertices transitions from the left to the right V1. On the right side of the figure, the median HWT of surviving features are displayed in graph plots projected into the visual field, with the same conventions as in Figure 8.

## Discussion

Between-subject prediction of the observed stimuli from intra-V1 correlation matrices (corresponding to their associated fMRI time-widows) was highly accurate for the four types of specific discriminations, both for stimulus level (Global vs. Local for both the ‘E’ and the ‘U’) and for letter shape (’E’ vs. ‘U’ for both Global and Local levels). This effect for simple classification was equivalent across all fMRI processing schemes. The simple effects were also found for simulated correlation matrices based on time-series modeling possible changes in retinal stimulation due to experimentally measured eye movements. In contrast, accurate cross-classification was only possible for level (ignoring shape) but not for shape (ignoring level). Furthermore, cross-classification of Level benefited from attenuation of the stimulus-evoked response, dropping to chance without deconvolution of stimulus-evoked response or when using beta-series. Interestingly, Level cross-classification was not due to eye-movement artifacts since this discrimination was at chance with the simulated correlation matrices. The weight maps for the level cross-classifications showed that inter-hemispheric links contributed most to the global cases and restricted intra-hemispheric links most to local cases of cross-classifications.

Highly accurate specific classifications are not surprising since they could arise from the highly distinctive pattern of retinotopic stimulation associated with each type of stimulus. Two facts suggest that they are related to multiple sources of fMRI activity. First, the high accuracies of specific classifications did not vary if the stimulus-evoked response was deconvolved or not. This suggests that the background fMRI activity could contain enough information for performing the discrimination but that the stimulus-evoked response either contains the same information as the background activity or is in synergy. Moreover, the simulation of gaze-shift-induced fMRI time series indicates that this is another potential source of the specific classifications. Therefore specific MVPA of intra-V1 correlations are not very informative.

Before discussing other results, we will address confounds due to differences in eye movements across stimulus types. Differences in eye movements in our control experiment were small, although somewhat larger stimuli than in the fMRI experiment were used. Larger stimuli would favor larger eye movements This contradicts one study (Sasaki et al., 2001), that found much larger horizontal eye movements when attending to global than to local stimuli. However, Sasaki et al. used very large global stimuli (about 30 DVA) and very small local stimuli (about 2.4 DVA), a global/local ratio of about 13. In contrast, both our global stimuli (under 5.3 DVA) and local letters (under 1.05 DVA) were small, global/local ratio of about 5. Here both levels were largely restricted to the fovea. Thus, our stimulus dimensions probably explain why gaze distributions were not very different between conditions.

Despite their small size, the variation in fixation hotspots between stimuli can apparently interact with stimulus retinotopic patterns inducing changes in the intra-V1 correlation matrices. Our simulated time series were generated from the eye movements measured in our control experiment with a Gabor pyramid model. Specific classifications based on these simulated time series were accurate. Although in this simulation, there was no noise, these results caution about the potential role that gaze-related artifacts can play in the analysis of intra-V1 connectivity.

The most interesting finding in this article was that within-individual cross-classification was accurate for abstract level (i.e., learning transfer for level occurred across letters identities). This contrasts with the failure in cross-classification for letter identity. Note that both cross-classifications depend on information not strictly tied to the retinotopic pattern of stimulation (Kaplan et al., 2015; Valdés-Sosa et al., 2020): they are more abstract than specific discriminations. On the one hand, level cross-classification benefitted from stimulus-evoked response deconvolution. This is congruent with the failure to cross-classify level based on beta-series (the fluctuation in strength over time of the stimulus-evoked response).

This implies that the correlations underlying the level of cross-classification are based on the fMRI background activity. In other words, MVPA of the stimulus-induced activations would tap into different patterns than the ones observed here. On the other hand, level cross-classification was not possible with the time series simulating differential effects of eye movements associated with stimulus types. Note that the fMRI is blurred over space and time by the hemodynamic function and anatomy (Lindquist et al., 2009); Prince et al., 2022), is contaminated by noise, and thus does not produce the crisp time series generated by our modeling. With real data, gaze-induced activity would be even less likely to enable level-cross classification.

Invariance for Level implies that the details of line configurations distinguishing ‘E’ and ‘U’ were abstracted away (Hübner & Volberg, 2005). Shifting attention towards the Local/Global is thought to occur by filtering out high/low spatial frequencies from the representation of the retinal input. (Flevaris et al., 2011, 2014). In a prior study by our group using activation-based MVPA (Valdés-Sosa et al., 2020), information about level (independent from shape) was found in the scene-selective cortex (medial ventral occipitotemporal and middle occipital areas). This area could be a source of feedback contributing to the intra-V1 network effect of level. When processing Navon figures, the control of spatial scale is essential (Flevaris & Robertson, 2016).

Decoding of shape that is tolerant to changes in size is present in the fMRI activations of higher-order-visual areas such as LOC (Grill-Spector, 2003). In our previous study with the same data used here, information about shape invariant to level was found in activations from these object-selective cortices (Valdés-Sosa et al., 2020). The reason that cross-classification for letters failed is unclear, but there are several hypotheses to explore. One explanation is that size-invariant representations do not influence intra-V1 networks through feedback. This is incongruent with the robust functional connections between the foveal region of V1 and LOC (e.g., Baldassano et al., 2016). Since the coupling between early and higher-order visual areas is selectively switched on or off according to task requirements (e.g., Al-aidroos et al., 2012), perhaps in our experiment level-invariant shape information feedback was not needed. However, these ideas are speculative until further experiments are carried out.

Edgewise HTW maps were very similar for the classifiers for level trained with data associated with different letters. This indicates (Tian & Zalesky, 2021) an intra-V1 functional connectivity topology t independent from stimulus shape. Edge HTW maps of the cross-classifiers for level had different topologies when the Global and Local ‘weighing in’. For weights indicating Global stimuli, many links crossed the vertical meridian in the lower visual field. For weights indicating Local stimuli, links were limited to single quadrants (especially the lower ones), with only a few crossings of the horizontal meridian. These results provide converging evidence for the previous work of our group using the same data but based on hypothesis testing using mass univariate edgewise and network-based statistics. We conclude that feedback influences are probable drivers of the intra-V1 networks since examination of these HTW shows that most discriminative links are longer than 4 DVA which exceeds the size of V1 receptive fields in the stimulated region.

The fact that the classifier weight maps described here and the significant links of our previous study both show stronger interhemispheric connections associated with Global stimuli in the lower visual field is consistent with evidence that Global visual perception is more accurate in the lower compared to the upper VF (Christman, 1993; Levine & McAnany, 2005; Previc, 1990). This advantage could be explained by greater sensitivity in the lower visual field to lower spatial frequency components (Niebauer & Christman, 1998), which are needed to extract Global shapes, including those in Navon figures (Flevaris & Robertson, 2016).

This study has several limitations. Higher-powered replications are needed, given the possibility of false positive results and inflated effect sizes in results from small samples (Button et al., 2013). Another limitation of this study is the lack of control of eye movements during the fMRI recording. Eye movements produce uncontrolled blurring of the retinotopic stimulus representation, which could weaken the correspondence of topologies across participants. The simulations we performed show a potential effect of the intra-V1 matrices that are best controlled by recording gaze positions during scanning. Here, a small range of stimulus shapes was used. More diverse stimuli (as used in our previous work, Iglesias-Fuster et al., 2014) would allow better testing of the mechanisms underlying the V1-connectivity patterns described here. These problems must be addressed in replication studies with more stimuli and participants, with the measurement of eye movements, and additionally examined with multivariate analysis of variance methods (Allefeld & Haynes, 2014).

As a final point, we underline that our results are congruent with studies using MVPA based on fMRI activations revealing that V1 carries visual information with a spatial range more extensive than its pRFs (Smith & Muckli, 2010; Williams et al., 2008). The pattern of connections between V1 cortical sites transcends the simple dependence on geodesic distance found during the resting state (Dawson et al., 2016). Purely feed-forward models do not easily explain these patterns of functional connections. Altogether, the data presented here supports the hypothesis of V1 as a “cognitive blackboard” V1 (Roelfsema & de Lange, 2016), which could play an essential role in cognitive processes such as perceptual organization, attention, and memory.

## Author contributions

Designed research: MOO, MVS, PVS, JPH Collected data: JPH, JIF Contributed analytic tools: MOO, MVS, PVS, DM Analyzed data: MOO, MVS Wrote the paper: MOO, MVS, PVS

## Acknowledgements

This work was supported by the VLIR-UOS project “A Cuban National School of Neurotechnology for Cognitive Aging”, and the National Fund for Science and Innovation of Cuba.

